# Selective effects of temperature on body mass depend on trophic interactions and network position

**DOI:** 10.1101/233742

**Authors:** Avril Weinbach, Korinna T. Allhoff, Elisa Thébault, Francois Massol, Nicolas Loeuille

## Abstract

Body mass is a key trait constraining interspecific interactions in food webs through changes in metabolic requirements. Because climate warming affects metabolic rates, it creates direct selective effects on body mass. Many empirical studies suggest that body mass decreases under warming, although important exceptions have been noted. We first analyze the evolution of body mass in a simple consumer-resource model to provide conditions under which a body mass increase or decrease may be expected. We then extend our model to a multi-trophic food web context that allows for the coevolution of body mass and of feeding preferences. We focus here on how the trophic position of a consumer influences its evolutionary response to warming under different scenarios for the temperature dependence of attack rates. We observe that body masses can remain constant or increase with temperature when attack rates are constant or increasing with temperature, while body mass reductions in response to warming are only expected when attack rates have a thermal optimum and populations are initially locally adapted. We also found that body masses at lower trophic levels vary less under warming than body masses at higher trophic levels, which may be explained by decreasing levels of stabilizing selection along food chains.

## Introduction

Accumulating evidence suggests that current global change, and in particular climate warming, affects the evolution of body masses. Many researchers regard decreases in body mass as one of the “universal responses” to warming, next to range shifts and changes in life-history traits (Brose et al., 2012; Daufresne et al., 2009; Emmrich et al., 2014; Sheridan and Bickford, 2011). Such downsizing is more pronounced for aquatic compared to terrestrial species (Forster et al., 2012), but it has been found for systems as diverse as phytoplankton (Yvon-Durocher et al., 2011), carnivorous mammals (Yom-Tov et al., 2010), fishes (Edeline et al., 2013), amphibians (Reading, 2007) or birds (Yom-Tov et al., 2006).

Body mass is considered to be a key ecological trait largely defining ecological rates and life history traits (Peters, 1986; Woodward et al., 2005). It constrains average home range size (Lindstedt et al., 1986), life spans and metabolic requirements (Brown et al., 2004) and also affects species interactions, e.g. when predators favor species in a given mass window (Brose et al., 2006b). Such allometric relationships have been extensively studied during the past decades and are known to enhance stability in complex food webs (Brose et al., 2006b). Body mass evolution can therefore affect individual metabolism and demography, as well as multi-species interactions, with important consequences for the structure and functioning of ecosystems (Loeuille and Loreau, 2006) and their ability to provide essential services (Ohlberger, 2013; Woodward et al., 2005). It is thus an urgent challenge to understand how increasing temperatures affect body mass evolution and the responses of complex ecosystems incurred by such eco-evolutionary feedbacks.

Warming-induced responses of average body mass of homeotherms are often explained in terms of physiological or metabolic constraints (Brown et al., 2004; Gillooly et al., 2001). Consider the classic Bergmann’s rule, which describes a geographical pattern where species of smaller body mass are typically found in warmer environments. Bergmann explained the observed pattern with a higher surface-to-volume ratio that allows for increased heat radiation per unit body mass (Bergmann, 1848). However, not all of the empirical studies agree with such variations (e.g. in insects, Shelomi 2012). A number of cases in which body mass shows instead a variable response, or even an increase, are summarized in the review by Gardner et al. 2011. Such deviations can neither be explained by Bergmann’s surface-to-volume argument, nor by metabolic constraints.

While not denying that variations in body mass are partly driven by individual scale constraints (metabolic or physiological), such variations also largely alter the ecological context for the considered population. It may lead to mass-dependent feedbacks at the population level, as well as changes in competitive and predatory interactions in the community (Ohlberger, 2013). Body mass is therefore likely selected not only by direct metabolic or energetic requirements, but also by changes in the network context. For example, Yom-Tov and Yom-Tov (2005) show that the body mass of Palearctic shrews in Alaska increased during the second half of the twentieth century in response to warming, contradicting the prediction of Bergmann’s rule. They proposed that such increases may be explained by a higher food supply, resulting from improved weather conditions for the shrew’s prey in milder winters. This study highlights that the evolutionary response of a target species to warming can depend on its trophic interactions with other species.

It is important to address such a dynamic community context, since empirical evidence suggests that global environmental changes exert pervasive impacts on both antagonistic and mutualistic species interactions, leading to major changes in network composition and ecosystem processes (Tylianakis et al., 2008). Biotic interactions and feedback processes can lead to highly complex, nonlinear and sometimes abrupt responses to climate change (Walther, 2010). A synthesis of species ecology, species evolution and biotic interactions is thus needed to generate reliable predictions of species responses to changing environments, but these fields have so far mostly been studied seperately (Lavergne et al., 2010). In this context, it is not yet clear in which cases the network context will accelerate, hamper or even counteract selection for smaller body mass due to metabolic or energetic constraints.

Trophic interactions between species shape selective pressures on body mass among species and trophic levels because predator/prey body mass ratios are constrained by the physical ability of predators to ingest prey of a certain mass. Moreover, these interactions are at the same time temperature-dependent (Englund et al., 2011; Rall et al., 2012). The most common ways of dealing with temperature-dependent (trophic) interaction rates is to either assume temperature dependency is weak enough to be ignored completely, or that trophic interactions increase following the Arrhenius equation (as for example done in Binzer et al. 2016; Fussmann et al. 2007; Vasseur and McCann 2005). The latter can be a good approximation within a certain thermal window, but it neglects additional processes occurring at higher or lower temperature regimes (see for example Peck 2016; Portner and Knust 2007; Tansey and Brock 1972; West and Post 2016). For instance, Sinervo et al. 2010 found that a whole group of lizard species can be so physiologically stressed by warming that they may not maintain any efficient activity, contradicting the prediction of monotonically increasing attack rates with temperature. A hump-shaped relationship between temperature and attack rates is thus more realistic when considering wide temperature ranges and has indeed been found in a large meta-analysis by Englund et al. 2011.

Several theoretical studies based on allometric relationships have already investigated the impact of warming on interaction networks. For example, it has been shown that warming stabilizes predator-prey dynamics at the risk of predator extinction (Fussmann et al., 2014), and strongly decreases the diversity of mass-structured predator-prey networks (Binzer et al., 2016). These studies only use the Arrhenius approach for attack rates and do not consider evolutionary dynamics. There is thus a lack of studies exploring how the relationship between temperature and attack rates affect food web evolution on broad thermal scales.

The first goal of our study is to understand how predator-prey interactions may interfere with metabolic and energetic constraints in shaping the evolutionary response of body mass to warming climate. More precisely, we ask how the temperature dependence of the attack rates can lead to increasing or decreasing body masses with temperature. We tackle this first question by focusing on a simple consumer-resource model that accounts for the evolution of the consumer body mass in addition to ecological dynamics. The temperature dependency of the attack rate is included via three different scenarios, namely a null hypothesis of temperature-independent attack rates, the Arrhenius approach and a hump-shaped relation with temperature, reflecting the underlying complexity of the foraging and ingestion processes. We use the adaptive dynamics framework (Dieckmann and Law, 1996; Geritz et al., 1998; Metz et al., 1992) to investigate consumer body mass evolution. The simplicity of the model allows us to obtain exact analytical solutions for variations in the selected body mass and conditions of consumer-resource coexistence in this eco-evolutionary context.

The second goal of the study is to understand how selection on body mass changes depending on the position of the evolving species in the food web. We therefore study variations in body mass at different trophic levels using a large community evolution model (for a review on such models see Brannstrom et al. 2012; Drossel and McKane 2005; Fussmann et al. 2007). The model produces, via numerical simulations, multi-trophic networks that emerge and evolve in a self-organized, temperature-dependent manner. This temperature dependence is implemented in exactly the same way as in the simplified consumer-resource model, allowing us to assess which patterns observed in the simple model hold across trophic levels in complex, multi-species communities.

## Models and Methods

We use a two-step approach to study the impact of warming on body mass evolution. A simple consumer-resource model is used to study the impact of temperature on consumer body mass evolution. This two-species model is a simplification of a more complex, multi-trophic community evolution model, which is used to study how body mass response depends on trophic position in complex networks. Our two models are based on the same ecological assumptions, but differ in their treatment of evolutionary dynamics, as explained in the following. In the following parts, asterisks (*) indicate the ecological equilibrium, and tilde (~) the evolutionary one. Whenever mentioned, log corresponds to the decimal logarithm.

### Ecological Processes

In both models, a consumer morph *i* is characterized by its average adult body mass *x_i_* (measured in kg) and its preferred prey body mass *f_i_*. These traits determine the feeding interactions, as illustrated in fig. 1. The rate of change of a consumer biomass density *B_i_* (measured in kg per m^2^) is given by:

**Figure 1:**
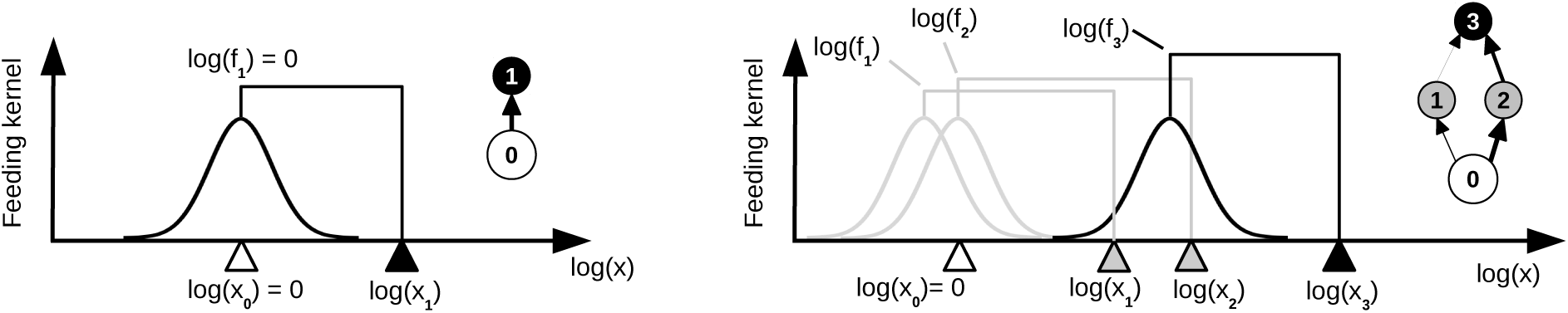
Illustration of the ecological rules in the two versions of the model. Left: A simple consumer-resource model. The consumer (black triangle) is characterized by its body mass *x*_1_ and the center of its feeding range *f*_1_. The Gaussian function (black curve) describes its attack rate kernel on potential prey, as explained in equation (2). The consumer thus feeds on the external resource (white triangle) with its maximum attack rate. Only the body mass *x*_1_ can evolve. Right: A more complex, multi-trophic model. Shown is a snapshot with three consumer morphs. Morph 3 (black triangle) feeds on morph 2 and 1 (gray triangles) with a high, resp. low, attack rate. Morph 1 and 2 are consumers of the external resource (white triangle). Note that both the body masses and the feeding centers can evolve, meaning that the network structures generated by this model are dynamic and typically more complex than this snapshot.

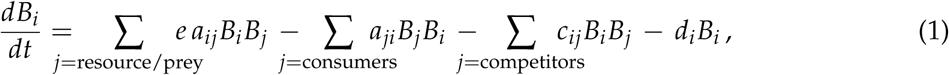

where *e* is the conversion efficiency, *a_ij_* is the mass-specific consumption rate at which morph *i* consumes morph *j*, *C_ij_* describes interference competition among consumers *i* and *j*, and *d_i_* is the respiration and mortality loss rate of consumer *i*. All ecological parameters are summarized in Table 1.

**Table 1:** A summary of all ecological model parameters both for the simplified consumer-resource model and for the more complex multi-trophic community evolution model. The models differ in the evolutionary rules, as explained in the methods section. ^*a*^ Values most commonly used in the analyses, and referred to as the typical values. ^*b*^ Values based on the work by Yodzis and Innes (1992). ^*c*^ Following the results of Gillooly et al. 2001 and Brown et al. 2004. ^*d*^ Values based on work from Binzer et al. 2016; Fussmann et al. 2014; Rall et al. 2012. ^*e*^ These parameters are chosen in a way that the attack rate is close to zero in case of *T* = 273K or *T* = 313K and maximal for *T_opt_* = 291,64K.

| Definition | Value(s)<sup>a</sup> | Unity |
| --- | --- | --- |
| $e$: Consumption efficiency, see (1) and (5) | 0.85<sup>b</sup> | [1] |
| $s$: Standard deviation of the feeding niche, see (2) | 0.25 | [log(kg)] |
| $c$: Scaling factor for interference competition, see (3) | 0.15 | $\left[\frac{\text{m}^2}{\text{s} \cdot \text{kg}}\right]$ |
| $r$: Resource growth rate, see (7) | 1 | $\left[\frac{1}{\text{s}}\right]$ |
| $K$: Resource carrying capacity, see (8) | 10 | $\left[\frac{\text{kg}}{\text{m}^2}\right]$ |
| $d$: Scaling factor for consumer loss rate, see (9) | 0.3<sup>b</sup> | $\left[\frac{1}{\text{s}}\right]$ |
| $E_a$: Activation energy used in (7), (8) and (9) | 0.65<sup>c</sup> | eV |
| $E'_a$: Activation energy used for $a_{ij}$ in (11) | 0.35<sup>d</sup> | eV |
| $T_{ref}$: Reference temperature used in (7) - (9), (11) and (12) | 293 | K |
| $T$: Local temperature used in (7) - (9), (11) and (12) | 273-313 | K |
| $k$: Boltzmann constant used in (7) - (9), (11) and (12) | $8.617 \cdot 10^{-5}$ | $\frac{\text{eV}}{\text{K}}$ |
| $b$: coefficient of the hump-shaped temperature dependency used in (12) | $b^e = -0.256$ | [1] |
| $q$: coefficient of the hump-shaped temperature dependency used in (12) | $q^e = -0.691$ | [1] |

Body mass and temperature dependence can affect the consumer loss rate, *d_i_* = *d_i_*(*x_i_, T*), and the consumption rates, *a_ij_* = *a_ij_* (*x_i_*, *x_j_*, *f_i_*, *T*). The different scenarios of temperature dependence are explained below. For a given temperature *T*, we assume that the consumer loss rate is constrained by body mass (Brown et al., 2004), 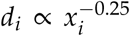 and that the consumption rate is a product of a metabolic scaling factor and a Gaussian feeding kernel, 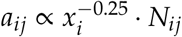. The feeding kernel (also illustrated in fig. 1) is defined as:

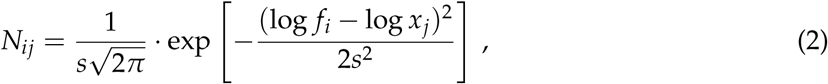

where the standard deviation s describes the degree of generalism of the consumer.

Following limiting similarity theory (Macarthur and Levins, 1967), we propose that competition increases when morphs have similar feeding niches. Their similarity is measured via the overlap of feeding kernels, *I_il_* = ∫ [*N_ij_* · *N_lj_*]*dw_j_*, with *w_j_* = log *x_j_*. We therefore write competition interaction as:

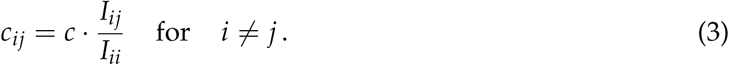

The model parameter *c* sets the overall competition strength in the system.

Energy input into the system is provided by an external resource with body mass *x*_0_ = 1. The rate of change of its biomass density *B*_0_ is given by

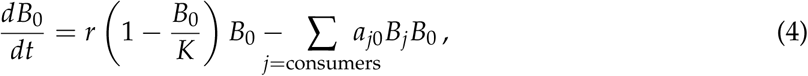

where *r* = *r*(*T*) and *K* = *K*(*T*) represent the temperature-dependent resource growth rate and carrying capacity, respectively.

### Evolutionary processes

In our first model, we consider a simple two-species system consisting only of the external resource and a single consumer population. Only the consumer body mass *x*_1_ is evolving, whereas its feeding center is fixed and matches the resource body mass, *f*_1_ = *x*_0_ = 1. In this simplified case, the population dynamics become:

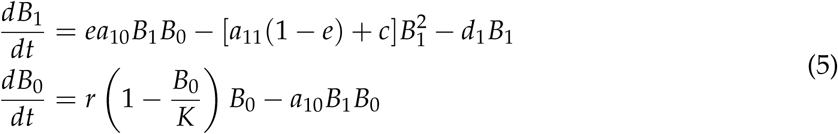

We follow the evolution of the consumer body mass *x*_1_ using the adaptive dynamics framework (Dieckmann and Law, 1996; Geritz et al., 1998; Metz et al., 1992). It assumes that evolution occurs via small mutation steps and that the system reaches the ecological equilibrium in between two mutations. Within this framework, the evolutionary dynamics of *x*_1_ is described by the canonical equation of adaptive dynamics (Dieckmann and Law, 1996):

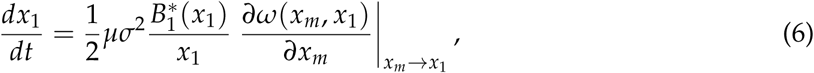

where 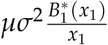 embodies the phenotypic variability on which selection can act, with *μ* the per individual rate of mutation, *σ^2^* the variance of the phenotypic effect of the mutation and 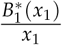 the density of the resident consumer population at equilibrium. The last term embodies variations in the fitness landscape around the resident value, thereby the effects of natural selection. The mutant is assumed to have a slightly different body mass value (*x_m_*) compared to a resident population fixing the ecological community (Metz et al., 1992). Because the part of the equation embodying the phenotypic variability is always positive, only the sign of the fitness gradient constrains the direction of trait variation. If the gradient is positive (resp. negative) then a higher (resp. lower) value of body mass is selected. In the results section, we use equation (6) to determine the position of evolutionary singularities and associated evolutionary dynamics.

In our second model, we consider not only the evolution of body masses, but also of feeding preferences, and we relax the strict assumptions of small mutation steps and separate ecological and evolutionary time scales. Such modifications facilitate the emergence of higher trophic levels and complex food webs. Each numerical simulation starts with a single consumer morph with body mass *x*_1_ = 100 and feeding center *f*_1_ = 1, which is thus feeding on the external resource with its maximum attack rate. The initial biomass densities are *B*_0_ = *K* = 10 for the resource and *B*_1_ = *∈* for the ancestor morph, where *e* is the consumer extinction threshold. Every 100 model time steps, we introduce a new morph into our network, which makes a total number of 5 · 10^6^ morph additions during a simulation runtime of 5 · 10^8^ time units. One of the extant morphs (but not the external resource) is chosen randomly as “parent” morph *i* for a “mutant” morph *j*. The mutation probability for a given morph is proportional to its individual density, meaning that morphs with small body masses on low trophic levels, which have the highest individual densities, have the potential to evolve fastest. The initial biomass density of mutants is taken from the parent morph and set to *B_j_* = *∈*. The logarithm of the mutant’s body mass log(*x_j_*) is randomly chosen from a normal distribution around the logarithm of the parent’s body mass log(*x_i_*), with a standard deviation of 0.25. The same rule applies also for the logarithm of the mutant’s feeding center log(*f_j_*). Parent and mutant morphs have thus typically similar traits, but bigger mutation steps can occasionally occur.

Whether or not a new mutant population is viable in a given environment created by the other morphs is determined by the population dynamics in equations (1-4). The resulting food web structures are not static, but emerge and evolve in a self-organized manner. Related community evolution models have been shown to produce complex networks with properties very similar to available empirical food web data (Allhoff and Drossel, 2013; Allhoff et al., 2015; Loeuille and Loreau, 2005). More details on the model that we use here, including exemplary simulation runs, can be found in the online appendix C.

### Temperature dependence

For both models, we include temperature dependence into the resource growth rate and carrying capacity, as well as into the consumer respiration and mortality loss rates:

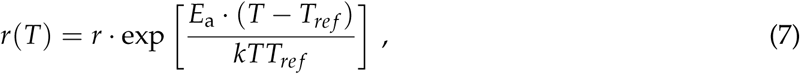

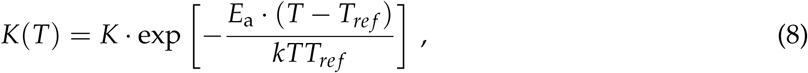

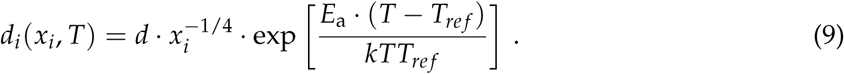

Here, *E_a_* is the activation energy, *T* is the local temperature, *T_ref_* is the reference temperature and *k* is the Boltzmann constant. *r*, *K* and *d* are scaling factors (see Table 1). Using this Arrhenius approach in order to include the temperature dependency into the resource growth rate *r* and into the respiration and mortality loss rate *d_i_* is consistent with previous studies by Gilbert et al. 2014 and Vasseur and McCann (2005). Our approach for the temperature dependency of the carrying capacity *K* is motivated by previous work from Binzer et al. 2012 and Meehan (2006), and consistent with the empirical observations reported in the analysis by Fussmann et al. 2014. For simplicity, we chose identical activation energies in all three cases.

We compare three different scenarios linking temperature with feeding interactions. Empirical data on attack rates reveal that the relationship with temperature is weak compared to the influence of temperature on parameters reported above (Fussmann et al., 2014; Rall et al., 2012, 2010). Thus, as a first approximation, we consider attack rates to be independent from temperature (scenario (a)), which serves as a null model:

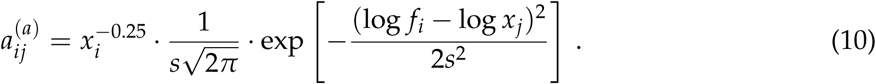

Scenario (b) assumes that temperature dependence of attack rates follow the Arrhenius equation (Binzer et al., 2016; Fussmann et al., 2007; Vasseur and McCann, 2005):

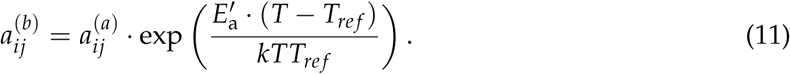

Following Rall et al. 2010, Rall et al. 2012 and Fussmann et al. 2014, we use a relatively low value for the activation energy 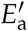 compared to the activation energy for the resource parameters and consumer loss rates, *E_a_*.

As explained in the introduction, continuous increases in attack rates under warming (as in scenario (b)) is only valid within a certain temperature range and is not suitable to describe temperature dependencies above the thermal optimum. In scenario (c), we therefore follow the results from Englund et al. 2011, and assume a modal relationship with a peak of attack rates at an optimal temperature (*T_opt_* = 291,64 K):

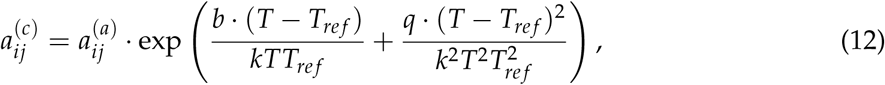

### Data collection

For both models and all three scenarios of temperature-dependent attack rates, we investigate how the consumer body mass(es), as well as the distribution of biomasses respond to warming. The simplified consumer-resource model can partly be treated analytically. We first study the ecological dynamics of the system, investigating the conditions for consumer-resource coexistence at a given temperature and for a given consumer body mass *x*_1_. We then study the evolution of consumer body mass using adaptive dynamics (Dieckmann and Law, 1996; Geritz et al., 1998; Metz et al., 1992). All analytical results from the consumer-resource model are corroborated by numerical simulations. Whenever adaptive dynamics equations are analytically intractable, graphical exploration of the possible evolutionary outcomes are made using pairwise invasibility plots (PIPs) (Geritz et al., 1998). Such PIPs visualize the invasion success of potential mutant populations given resident populations via the sign of their relative fitness. They allow us to investigate whether evolutionary singularities occur and whether they are convergent and invasible.

The multi-trophic community evolution model is analyzed via numerical simulations only. Each simulation allows the emergence of a network. After an initial period of diversification, the network size and structure stays approximately constant and fluctuates around a temperature-dependent average. We are particularly interested in the fluctuations in network structure and biomass flow after this initial build-up, and therefore deliberately exclude the first 5 · 10^7^ time units from the analysis. Throughout the simulations, we calculate the trophic positions of all morphs via the average, flow-based trophic position of their prey plus one. We round these trophic positions to integer values in order to group the morphs into distinct trophic levels. We then determine the average body mass and the total biomass of all morphs found at each trophic level and finally calculate the time average of these measures. More detailed information on this procedure, including exemplary simulation runs, can be found in the online appendix C.

## Results

### Ecological equilibria

We start with the analytical investigation of the simplified consumer-resource model. System (5) leads to three possible ecological equilibria. Two of those are trivial, because either the consumer 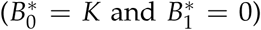 or both species 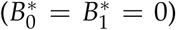 go extinct. A third equilibrium allows for the coexistence of both species and is therefore of particular interest. The equilibrium biomass of resource and consumer are given by:

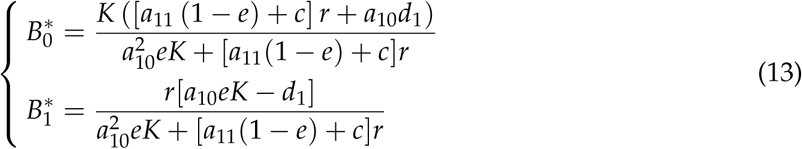

Note that this equilibrium depends on all parameters and in particular on the consumer body mass *x*_1_ that affects the metabolic scaling of loss (*d*_1_) and attack rates (*a*_10_ and *a*_11_). Feasibility and stability conditions required to maintain the coexistence are detailed in the online appendix A.

### Impact of temperature-dependent attack rates on body mass evolution and biomass densities

In this part, the parameters depending on body mass are rewritten as functions of this trait (e.g. *a*_10_ becomes *a*(*x*_1_,*x*_0_)). As explained in the methods section, evolution of the consumer body mass is determined by the fitness gradient (see equation(6)). The relative fitness *ω*(*x_m_*, *x*_1_) of mutant consumer *m* (with biomass density *B_m_* and body mass *x_m_*) given the resident consumer 1 (with biomass density 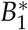 and body mass *x*_1_) is determined by the mutant population growth rate when rare and the resident being at ecological equilibrium:

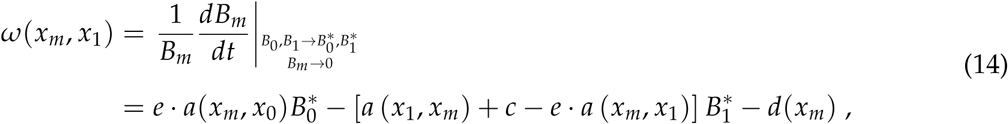

where *a*(*x*_1_, *x_m_*) describes the predation rate of the mutant by the resident, *a*(*x_m_*, *x*_1_) is the preda tion rate of mutants on residents, and *d*(*x_m_*) is the mutant death rate.

The fitness function can be used to determine evolutionary singular strategies, which occur when trait variations are null. This correspond to the roots of equation (6). Because other parts of the equation (6) are strictly positive, evolutionary singularities correspond to the roots of the fitness gradient 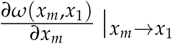. Computing this fitness gradient yields the following evolutionary singularities:

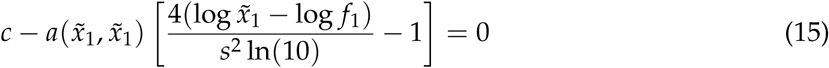

where the tilde indicates singular strategies. The complete proof of this result, as well as numerical conditions for non-invasibility (i.e., conditions under which such strategies cannot be invaded by any nearby mutant) differentiated from this equation, can be found in the online appendix B.

While equation does not allow an explicit expression for selected consumer body mass 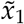 (15), it reveals which parameters affect its evolution under warming. More precisely, 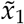 clearly depends on competition and predation rates, but not on resource parameters or consumer loss rates. Effects of temperature changes can then be directly analyzed. Taking the derivative of (15) with respect to T shows that variations the singular strategy 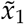 are positively related to variations of the consumer attack rate with temperature (none in scenario (a), Arrhenius-shaped in scenario (b) and hump-shaped in scenario (c)), for the vast majority of parameter sets. The direction of body mass evolution in response to warming is thus entirely determined by the direction of the effect of temperature on consumer attack rate. A complete analysis can be found in the online appendix B.

Pairwise invasibility plots (PIP) (Geritz et al., 1998) corroborate these findings (fig. 2). For all three scenarios and for the whole temperature range considered in our study, we always find two singular strategies: a repellor and a continuously stable strategy (CSS). Consumers with a body mass close to the repellor will evolve away from it (as shown on panel C), whereas a population near a CSS will evolve towards the singular strategy and settle there (as shown in panels B and C). Consistent with our analysis of equation (15), the position of the CSS is temperature independent in scenario a). Panels D, E and F confirm that increasing temperature leads to continuously increasing consumer body mass under scenario (b). Scenario (c) first leads to an increase (panel G and H) then to a decrease (panel I) of consumer body mass.

**Figure 2:**
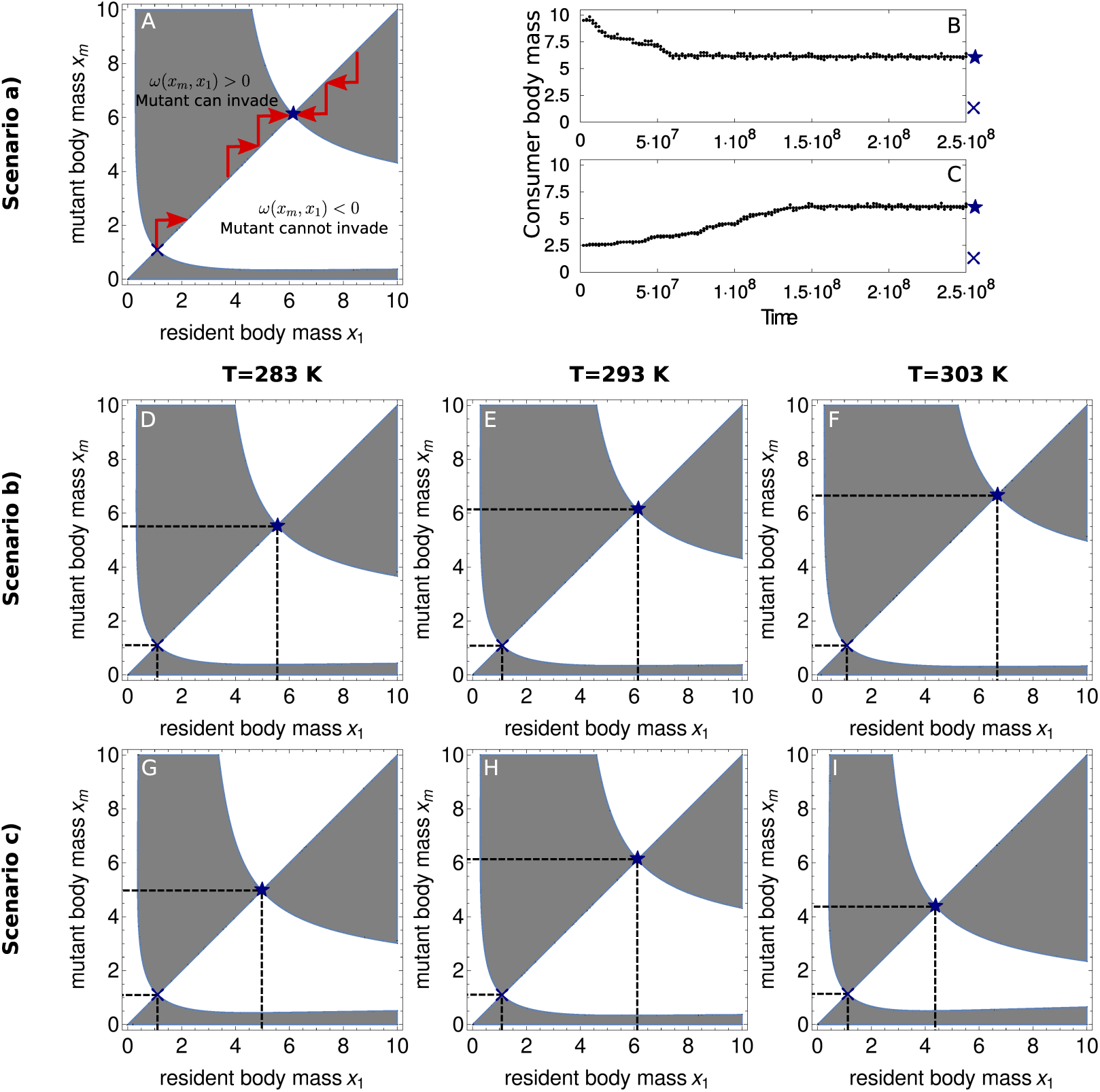
A, D to I: Pairwise invasibility plots (PIP) representing the potential invasibility of a rare mutant with body mass *x_m_* within a population of resident consumers at the ecological equilibrium with body mass *x_1_*. In A, *ω (*x*_m_*, *x*_1_) is the relative fitness of a mutant with trait value *x_m_* compared to a resident with trait value *x*_1_. Evolutionary trajectories are represented by red arrows. The system has two evolutionary singular strategies, one non-convergent and invasible (the blue cross, called a repellor) and one convergent and non-invasible (the blue star, called a continuously stable strategy or CSS). B and C: Simulations of consumer body mass evolution under scenario (a) starting at *x*_1_ = 10 (B) and *x*_1_ = 2.5 (C).

Fig. 3 further illustrates the variations in body mass due to temperature, as well as their implication for distribution of biomasses. Panels A, C and E show the variations in the selected (CSS) body mass for a wide range of temperature and for each of the three scenarios tested. Increasing attack rates (as in scenario (b) or scenario (c) when below the thermal optimum) lead to increasing body masses under warming. Decreasing attack rates (as for high temperature in scenario (c)), in turn result in decreasing body masses under warming.

**Figure 3:**
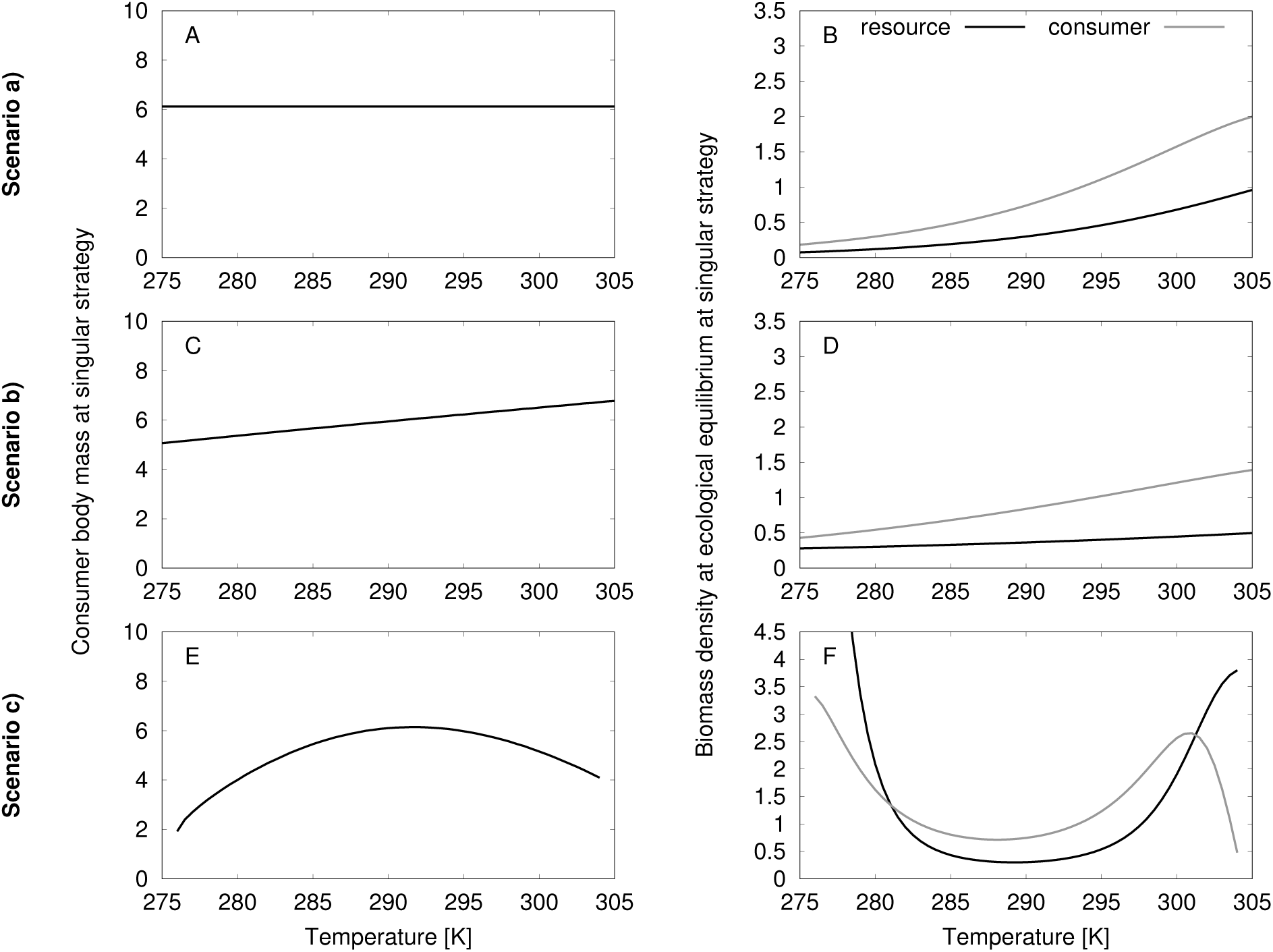
Analysis of the consumer-resource model with three scenarios of temperature-dependent interaction rates. A, C, E: Impact of temperature on final consumer body mass 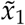. B, D, F: Impact of temperature on biomass densities 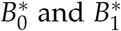. Parameters values are the typical one specified in Tab.1.

The final distribution of biomasses at the end of the eco-evolutionary process also depends on the considered scenario for the consumer attack rate (fig. 3B, D, and F). In scenario (a), both 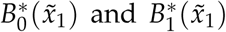 increase with temperature (panel B), as a direct consequence of increasing resource growth rates (equation (7)). The decline in carrying capacity under warming (equation (8)) gets more important at even higher temperatures, and eventually leads to consumer extinction, but plays only a minor role in the temperature window considered here. Such a pattern is also consistent with relaxed top-down controls (Oksanen et al., 1981), as the consumer suffers increasing loss rates (equation (9)). Warming eventually leads to a decreasing consumer-resource biomass ratio.

A similar pattern of increasing biomass densities, but with an increasing consumer-resource biomass ratio, emerges from scenario (b), because increasing attack rates (equation (11)) partly compensate increased loss rates (panel D). In contrast, the pattern emerging from scenario (c) is more complex (panel F). Coexistence of resource and consumer is only possible within a certain temperature window. At high temperature, the reduced attack rate does not allow the maintenance of the consumer (equation (12)). Maximum attack rates at intermediate temperatures lead to resource depletion, explaining the U-shape of the resource biomass density with temperature.

### Impact of trophic positions on body mass evolution and biomass densities

The multi-trophic community evolution model allows us to check whether the results that we obtain from the simplified consumer-resource model, hold across trophic levels in a more complex network context. We therefore explore whether and how the trophic position of a consumer affects its evolutionary response to warming. In general, we obtain results that are very similar to the results from the simplified consumer-resource model, as shown in panels A, C and E of fig. 4: Consumer body masses do not change in response to warming under scenario (a), increase under scenario (b) and show a hump-shaped response under scenario (c). However, we observe that the variation in body mass observed in scenario (b) and (c) is systematically larger at higher compared to lower trophic levels. Changing temperatures trigger evolutionary responses on all trophic levels, meaning that consumers not only react directly to changing temperatures, as in the simplified model version, but also to changing prey and/or predator body masses. Evolutionary changes in body masses thus cascade through the whole food web and trigger eco-evolutionary feedbacks. While lower levels are heavily constrained (selected body mass being a compromise of many effects, including body mass of both predators and prey), top levels evolve more freely (as their body mass is not constrained by any predator above). This results in decreased levels of stabilizing selection at higher trophic positions.

**Figure 4:**
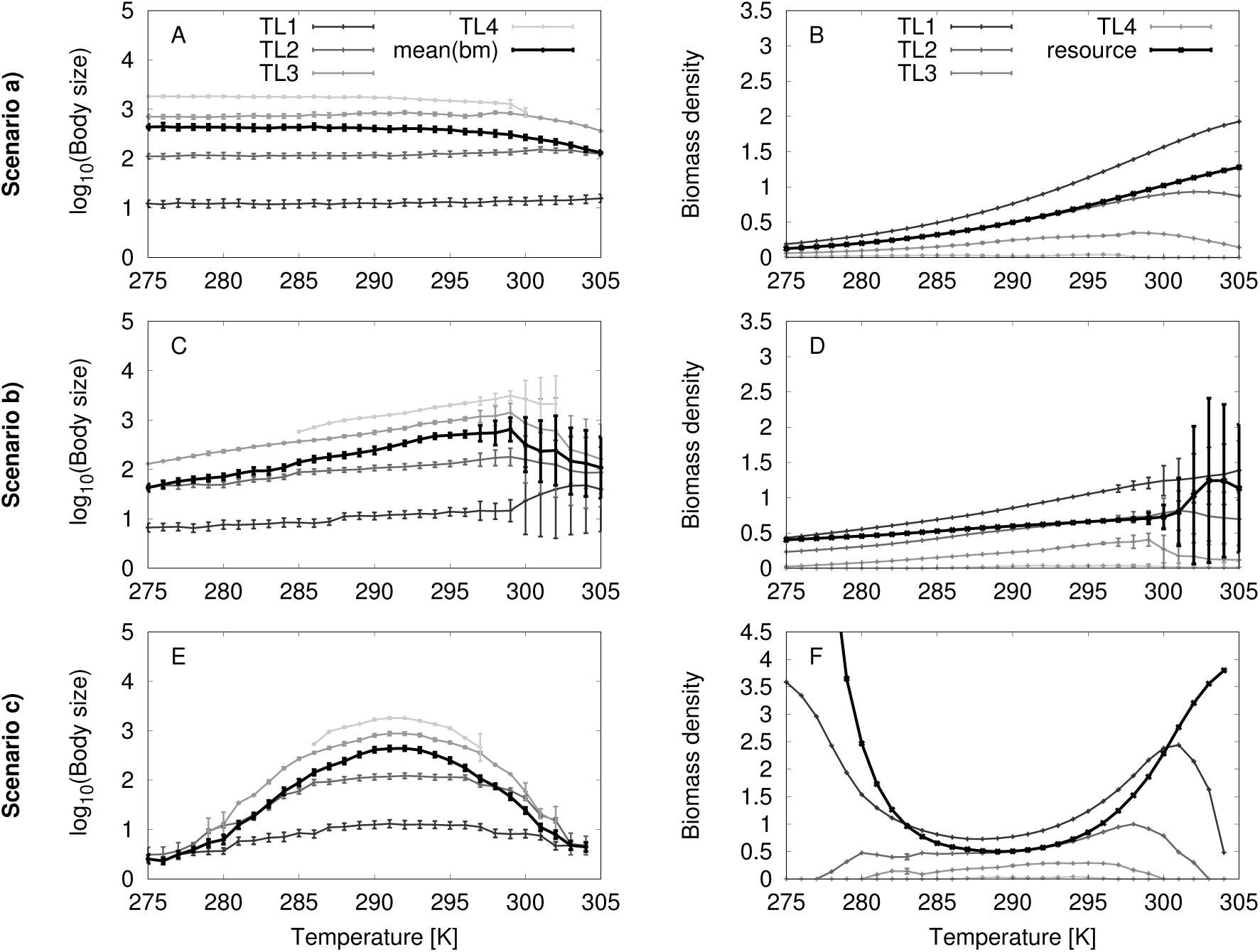
Same analysis as in fig. 3, but with the multi-trophic community evolution model. A, C, E: Body mass response to warming. Shown is the mean body mass of all consumer morphs in the network and of all consumers morphs within a given trophic level. Each data point represents an average over time and over 10 simulations runs. Error bars represent standard deviations describing the variation among simulations. The initial network build-up is not taken into account. B, D, F: Response of the biomass distribution across trophic levels to warming. Shown is the biomass density of the resource and the total biomass density of all consumer morphs within a given trophic level. Each data point represents again an average over time and over 10 simulations runs.

The response of biomass densities to warming is a straightforward generalization of the consumer-resource model, as shown in panel B, D and F of fig. 4. Warming in scenario (a) leads to an increase in resource biomass density, but decreasing consumer efficiency hampers the biomass flow from lower to higher trophic levels and therefore leads to starvation of top predators in hot temperature regimes. Scenario (b) leads again to a similar, but less pronounced pattern, reflecting the fact that increasing attack rates compensate for increasing consumer loss rates. Scenario (c) leads to resource depletion at intermediate temperatures, where attack rates are maximal. Increasing (or decreasing) temperature first leads to starvation of top predators, which temporally releases the next lower trophic level from predation pressure, and then these new top predators also go extinct, and so on, until consumer survival is impossible.

## Discussion

Our results reveal that evolution of body mass in response to warming can greatly depend on the temperature dependence of consumer attack rates and that such body mass changes cascade through the whole food web. We investigated three different scenarios: (a) Temperature-independent attack rates, (b) attack rates increasing with temperature, following the Arrhenius approach, and (c) a hump-shaped relation between temperature and attack rates. By comparing results obtained under these three different scenarios, we uncovered that body mass increase or decrease follow the variations of attack rates, throughout the network. Our approach considers both organism-level metabolic and ecological constraints linking temperature and body mass (i.e. temperature-dependent resource growth, respiration and attack rates, and temperature-dependent resource capacity), but does not account for developmental or cell-level metabolic rationales (such as those invoked by Arendt, 2007; Kozłowski et al., 2004; Perrin, 1995; van der Have and de Jong, 1996). Our approach thus assigns body mass variations to the result of ecological selective pressures acting directly at the organism mass level (Atkinson and Sibly, 1997; Daufresne et al., 2009; Hessen et al., 2013).

The simplified model suggests that temperature effects on body mass greatly depend on how temperature constrains attack rates. We find that scenario (a) and (b) lead respectively to no change and to an increase in consumer body mass under warming, whereas scenario (c) results in a hump-shaped relation of body masses with temperature. The decline in consumer body mass, as observed in numerous empirical studies (Brose et al., 2012; Daufresne et al., 2009; Emmrich et al., 2014; Sheridan and Bickford, 2011), thus only occurs under scenario (c), under the assumption that the consumer was initially adapted to thermal constraints and now displays decreasing attack rates with warming. The assumption of an increase in attack rates as a first response to warming (such as in scenario b) may be justified as consumers first need increased energy levels to face new metabolic requirements. During this first response we therefore expect, according to our results, an increase in selected body mass. However, empirical evidence shows that many species may already be limited in their daily activities, including foraging, as they have to spend time in refugia to prevent overheating (Sinervo et al., 2010). Such observations suggest that such species have passed their thermal optimum, so that attack rates decline, as in scenario c. We then expect, consistent with most reported empirical results, that declines in selected body mass should be expected for such species. In a world that has already warmed significantly, scenarios (a) or (b) might be limited in scope, so that no variation or increases in body mass may seldom be observed.

By allowing different outcomes in terms of body mass variation, our model helps to account for the important exceptions to the supposed universal rule of declining body masses with climate warming (Gardner et al., 2011). Empirical examples related to scenario (a), which show no body mass response to warming, are likely to be under-reported in the empirical literature, as negative results are more prone to self-censorship, and/or harder to publish. However, according to our model, we still expect frequent evolution to smaller body masses. Indeed, the assumption leading to such an outcome (locally adapted consumers, which suffer from decreased consumption efficiency when being forced to leave their thermal optimum under global warming), seems to be reasonable in many cases. This assumption of local adaptation even forms the cornerstone of so-called climate envelope models that take current species distributions as reflections of their niches to predict future species distributions under changing environmental conditions (Thomas et al., 2004). It is likely fulfilled for species with large populations and large spatial ranges, as selective pressures then have ample opportunities to act and allow local adaptation.

The mechanistic explanation for this direct link between attack rate and body mass variation is as follows: in the simplified consumer-resource model, the selected consumer body mass is constrained by two conflicting pressures. The first pressure is energetic and corresponds to the balance between feeding input and biomass loss terms. It favors rather small body masses. All of these terms indeed scale with body mass, except for competition, so that a smaller morph overall experiences less competition pressure. By contrast, the second pressure is due to cannibalism and favors big body masses that suffer less such additional mortality. Scenario (b) reinforces the strength of the second component (by increasing attack rates under warming) relative to the first force, so that the body mass increases with warming. The same argumentation also holds for scenario (c) when temperature is below the optimum. Above the optimum, the trends get reversed and lead to decreasing body masses.

The direct link between evolved body mass and the temperature-dependence of consumer attack rate is confirmed by our mathematical analysis of the simple model. Note however that it relies on the assumption that resource parameters and consumer loss rates have identical activation energies. Temperature dependencies may however largely vary among parameters, as for example suggested for the resource carrying capacity by Uszko et al. 2017 and O'Gorman et al. 2017. In this light, a complete understanding of the temperature dependencies remains an important question, that goes beyond the scope of this article. We can nevertheless assume that our first key result, which states that substantial decreases in body masses are only observed under scenario (c) as a consequence of decreased attack rates, is at least robust against choosing slightly different activation energies in (7), (8) and (9), because their impact would be overcompensated by the impact of considering or neglecting temperature-dependent attack rates. This interpretation is perfectly in line with the study by Edeline et al. 2013, who predict that warming-induced body downsizing emerges through both ‘direct’ (ecology-independent, e.g. thermal constraints on physiology) and ‘indirect’ (ecology-mediated, e.g. shifts in selection induced by species interactions) effects of temperature, but that ecology provides the overwhelming forces driving thermal clines in fish body mass.

Direct links between attack rate and selected body mass variations may be empirically tested in different ways. A first test could focus on selected examples of body mass responses confirming or contradicting the universal rule of declining body masses with temperature (Atkinson and Sibly, 1997). Targeted experiments on variation in performance and attack rates in response to temperature would allow to determine whether the considered species or populations are at a particular position of their thermal niches (see e.g. Dreisig, 1981; Englund et al., 2011; Fedorenko, 1975; Gresens et al., 1982; Mohaghegh et al., 2001; Thompson, 1978, for classic studies and a meta-analyses investigating interaction rates at various temperatures). We predict that an increase (resp. decrease) in body mass is related to consumers that were initially adapted to temperatures warmer (resp. colder) than their environment and therefore benefit (resp. suffer) from warming. We further predict that those cases where no body mass response is observed reflect virtually temperature-independent attack rates (as might be the case for small temperature changes around the consumer thermal optimum) or situations where the species had no evolutionary potential to follow the temperature change (e.g. because it is very rare, has a low genetic variability or long generation times compared to the pace of climate warming).

A second test could be based on experiments using organisms with a short generation time such as phytoplankton and zooplankton species, which have often been successfully used to investigate eco-evolutionary processes (e.g. Yoshida et al. 2003, Pantel et al. 2015). Growing several populations at different temperatures and confronting these pre-selected populations with different temperature regimes would allow a manipulation of the species position relative to its thermal optimum. Based on our scenario (c), we predict that the body masses of organisms selected at temperatures below (resp. above) the species' thermal optimum will increase (resp. decrease) with warming.

Interestingly, the qualitative variations of body masses observed in the simplified consumer-resource model hold in more complex, multi-trophic communities. We find, however, that the observed relative changes in body mass are larger at higher trophic levels compared to lower trophic levels. Although all consumers are directly affected by changes in temperature through their attack and death rates, as in the simplified model, most consumers also have to adapt to changes in prey and/or predator body mass. As a consequence, we observe that body mass changes cascade through the whole food web. Higher trophic level species undergo less stabilizing selection than those at lower trophic levels, because low trophic levels are constrained by the body masses of both resource and predator species, whereas top predators are not constrained by any predator and thus evolve more freely.

That temperature changes interact with trophic interactions to select body mass variations is in line with empirical observations. Gibert and DeLong (2014), who analyzed a large marine data set uncovered that prey mass selection depends on predator body mass, temperature and the interaction between the two. Our finding that sensitivity to warming increases with increasing trophic position is also in line with data from multi-trophic grassland communities (Voigt et al., 2003). While top predators show larger evolutionary responses of body masses in our model, they also are the first to go extinct under warming. This finding is again in line with several empirical observations (Beisner et al., 1997; Gibert and DeLong, 2014; Petchey et al., 1999) and can be explained by the low abundance at the top trophic levels. Increased respiration/death rates due to warming then lower the population growth rates of morphs that are already rare. Such extinctions are therefore coherent with classic works on top down controls (Oksanen et al., 1981) or with other works that link trophic length and energy availability (Pimm, 1982; Post, 2002).

While the flattening of trophic networks under warming agrees with previous results (Arim et al., 2007; Fussmann et al., 2014), most previous works rely on ecological processes only, ignoring the role of (eco-)evolutionary dynamics. One might for example imagine that those trophic levels that disappear in response to warming can re-emerge, meaning that evolution eventually “repairs” the damage that took place (Kokko et al., 2017), or that body mass evolution helps to maintain constant consumer-resource biomass ratios and buffer the community from extinctions in the first place (Osmond et al., 2017). Our model, relying on eco-evolutionary dynamics, can thus help clarifying this question. Evolution is clearly not sufficient to completely restore (or maintain) the network structure after (during) warming. Instead, we predict that warming will significantly change the food web structure not only through variations in population density due to changes in ecological dynamics, but also due to changes in selected body massed. A related study by Stegen et al. 2011 predicts diversity to increase with temperature if resource supply is temperature-dependent, whereas temperature-dependent consumer vital rates cause diversity to decrease with increasing temperature. Combining both thermal dependencies (similar to our scenario b) results in a unimodal temperature-diversity pattern. A more detailed analysis of how temperature shapes evolving food web structures is, at least to our knowledge, still lacking.

Our observation that consumer body mass evolution is more sensitive to warming at higher trophic levels, could be tested with empirical data, since body mass distributions are now widely measured in food webs (Brose et al., 2006a; Cohen et al., 2003; Petchey et al., 2008; Riede et al., 2010; Woodward et al., 2005). However, we are confronted with two major difficulties when using such data sets. First, we would need to have these distributions on many generations, which is challenging, especially since high trophic levels are typically occupied by long generation species. This first obstacle can be overcome by using palaeontological data (Willis and MacDonald, 2011) or indirect evidence, e.g. from a space-for-time substitution, with the usual caveat that latitudinal gradients correlate not only with temperature changes, but also with other environmental variables (eg, growth season duration) (Hessen et al., 2013).

Second, current systems are not only stressed by warming, but also by other changes, such as nutrient availability or habitat fragmentation, which are already known to affect food web complexity (Calcagno et al., 2011; Pillai et al., 2011; Post, 2002). These simultaneous changes can affect body mass evolution and food web dynamics in ways that are antagonistic or synergistic to warming effects. This second obstacle can only be addressed with controlled experiments, for example using mesocosm experiments. While such mesocosm experiments may have some limits in terms of representing natural network complexity (eg, aquatic mesocosm most often rely on phytoplankton-zooplankton, but may oversimplify diversity at upper trophic levels), they have provided important tests for ecological theories (see for example (Hulot et al., 2000) or (Downing and Leibold, 2002)) and are currently developed to understand the effects of global changes on complex system assemblages (Yvon-Durocher et al., 2011).

Based on our investigation, we see two important challenges for future research. The first challenge focuses on food web structure and stability, as already indicated above. A recent modeling approach by Binzer et al. 2016 led to the conclusion that the persistence and the connectance of complex, mass-structured predator-prey networks decrease with warming. It is, however, unclear whether these predictions hold when evolutionary dynamics, and in particular body mass evolution, is taken into account. We know that eco-evolutionary feedbacks can provoke surprises concerning species coexistence, through evolutionary rescue effects (Bell and Gonzalez, 2009; Ferriere and Legendre, 2013; Gonzalez et al., 2013) or evolutionary extinction debts (Norberg et al., 2012). In some cases, body mass evolution might help to maintain biodiversity by modifying consumer-resource mass ratios and thereby altering interaction strengths and energetic efficiencies (Osmond et al., 2017; Sentis et al., 2017), but the differential sensitivity of trophic levels to warming might also lead to community destabilization (Voigt et al., 2003). A change in network structure in response to warming also influences the functioning of the network and hence its ability to provide essential ecosystem services (Allhoff and Drossel, 2016). We therefore strongly suggest further research on the question how food web structures change in responses to warming, how evolution shapes these responses, and consequently what changes in the stability and functioning can be expected.

The second challenge addresses spatial aspects of food web eco-evolutionary dynamics. In our study, we assumed well-mixed populations in a homogeneous landscape and neglected any kind of spatial dynamics. It has, however, been predicted that gene flow and invasions have the potential to affect local adaptation and vice versa, depending on the relative timescale of spatial and evolutionary dynamics (Calcagno et al., 2017; Kirkpatrick and Barton, 1997; Norberg et al., 2012; Urban et al., 2012). It has also been predicted that spatial dynamics occurring between coupled habitats have the potential to change selection pressures in local food webs (Bolchoun et al., 2017). Evolutionary metacommunities, which integrate large community evolution models (Brannstrom et al., 2012) with the concept of metacommunities (Leibold et al., 2004; Pillai et al., 2011), might thus be key to generate a thorough understanding of evolutionary responses to global warming.

## Acknowledgements

This work was supported by the French National Research Agency (ANR) through project ARSENIC (grant no. 14-CE02-0012). AW was additionally supported by the École Normale Supérieure de Lyon. We thank Guillaume Chero, Alexandre Terrigeol and Elise Kerdoncuff, who performed simulations and robustness checks using a preliminary model version during their master internships. We also thank Youssef Yacine for helpful discussions and feedback on the manuscript.

## Author contribution statement

AW and KTA share the first authorship for this article. All authors participated in designing the models. AW performed the analytic calculations of the consumer-resource model, while KTA performed all numeric simulations of both models. All authors discussed the results. AW and KTA wrote the first draft of the manuscript. All authors then participated in the manuscript writing.

## Online Appendices

### A: Ecological analysis of the consumer-resource model

#### Method

Variations of the consumer and resource biomass densities with time (ecological dynamics) are described by equation (5) in the main article. Of special interest are conditions that determine the equilibrium of consumer and resource biomass densities, meaning the solutions of equation (5) equal to (0,0). The evolutionary analysis we undertake requires that the equilibrium is feasible and stable, so that we need to determine the parameters ranges for equilibrium feasibility and stability.

Feasibility simply means that biomasses are non negative. Studies usually consider positive or null biomasses, but because we focus here on species interactions, we consider strict positive biomasses for both species: 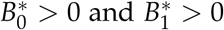.

Local stability is investigated by analyzing the associated Jacobian matrix (***J***). Stability requires: *det*(***J***) < 0 and *Tr*(***J***) > 0.

#### Results

Feasibility and stability conditions for the three scenarios are presented in table A1 below. We can show that in our case, if feasible, the equilibrium is always locally stable.

**Table A1:** Feasibility and stability conditions

|  | Feasibility<sup>a</sup> conditions at low K | Feasibility conditions at high K |
| --- | --- | --- |
| Scenario (a) | $0 < T < T_{lim} = \frac{T_{ref}E_a}{E_a - T_{ref}k \ln(\sqrt{\frac{a_{10}eK}{d}})}$<br>and $0 < K < \frac{d \exp(\frac{2E_a}{T_{ref}k})}{a_{10}e}$ | $0 < T \text{ and } K \geq \frac{d \exp(\frac{2E_a}{T_{ref}k})}{a_{10}e}$ |
| Scenario (b) | $0 < T < T_{lim} = \frac{T_{ref}(2E_a - E'_a)}{2E_a - E'_a + T_{ref}k \ln(\frac{a_{10}eK}{d})}$<br>and $0 < K < \frac{d}{a_{10}e}$ | $0 < T \text{ and } K \geq \frac{d}{a_{10}e}$ |
| Scenario (c)<sup>b</sup> | There is a lower and upper $T_{lim}$ | |

<sup>a</sup>: The stability conditions are always the same or less restrictive than the feasibility conditions.
Thus, when a system is feasible, it is always stable. <sup>b</sup>: Conditions for stability and feasibility are
given by the interval between the roots ( $T_{lim1}$ and $T_{lim2}$ ) of the following inequality:

### B: Evolutionary analysis of the consumer-resource model

#### Proof of the singular strategy formula

Adaptive dynamics are formalized by the canonical equation (Dieckmann and Law, 1996):

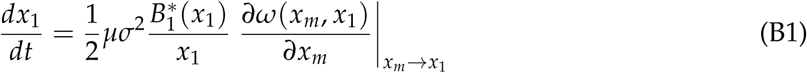

with all terms explained in the methods section of the main article. In this equation, the last term corresponds to the selection gradient, whose sign controls the evolutionary outcome (all the other terms are always positive and are therefore not influencing the direction of evolution of our focal trait *x*_1_). If the gradient is positive (resp. negative) then a bigger (resp. smaller) body mass is selected. Here the selection gradient is the derivative of the relative fitness of a rare mutant within a resident population. The selected trait is the body mass, with mutant body mass *x_m_* slightly different from that of the resident *x*_1_.

Calculation of the selection gradient requires the expression of the relative fitness function. In simple deterministic continuous-time models such as ours, relative fitness *ω* (*x_m_*, *x*_1_) of a mutant strategy *x_m_* is measured as the growth rate of a rare mutant population, given that the resident population is at its ecological equilibrium. It corresponds to equation (B2):

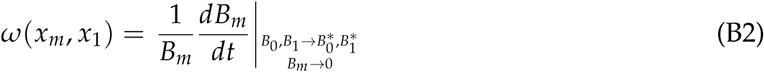

Let us consider interactions between the mutant and the resident in our present model, with the resource and resident consumer biomasses at ecological equilibrium and a rare mutant consumer. The growth rate is given by:

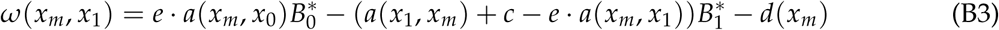

The first derivative of this fitness function gives the following selection gradient:

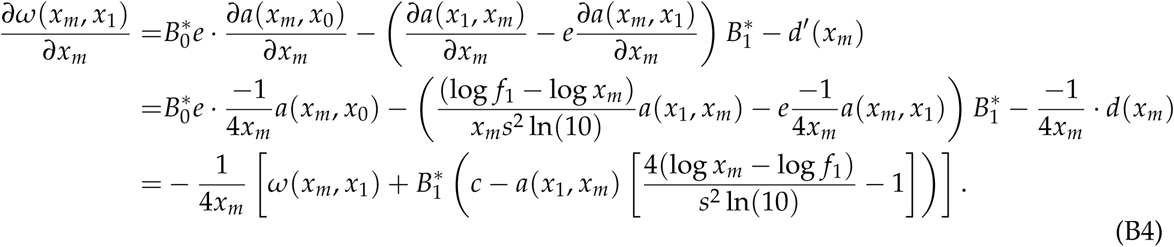

Evolutionary singularities occur at trait values which make the canonical equation (B1) equal to zero. Because other parts of equation (B1) are strictly positive, evolutionary singularities correspond to the roots of the selection gradient. Here the singular strategies 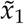 therefore correspond to the roots of equation 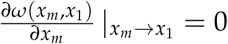. As mutations are small (*x_m_* goes to *x*_1_), and remembering that *ω* (*x*_1_, *x*_1_) = 0, equation B4 simplifies to:

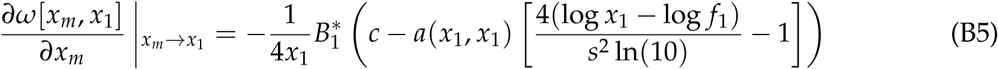

Remembering that 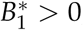, such singular strategies follow the equation:

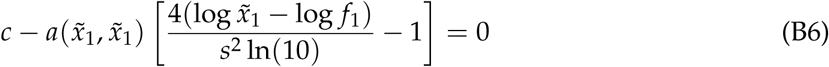

where the tilde indicates body mass values at the singular strategy.

#### Temperature dependence of the singular strategy

In order to study the variation of 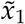 with the temperature *T*, we can differentiate (B6). Consider now 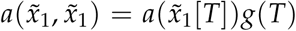, with *g*(*T*) the impact of temperature on the attack rate. *g*(*T*) = 1 in scenario (a), 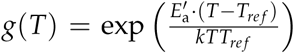in scenario (b) and 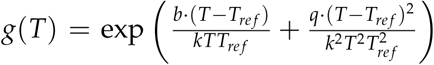 in scenario (c), with all the terms defined in table 1. Finally let us set 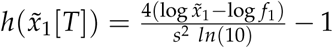.

With these new notations, (B6) becomes:

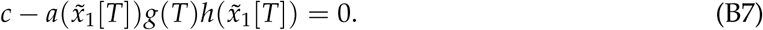

To understand the impact of temperature on the consumer body mass at the singular strategy 531 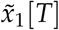, we differentiate (B7) with respect to temperature *T*:

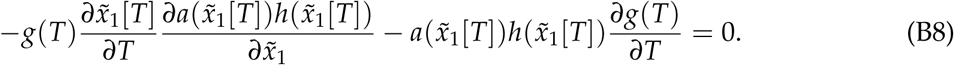

Then the sensitivity of 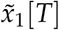 to changes in T is equal to:

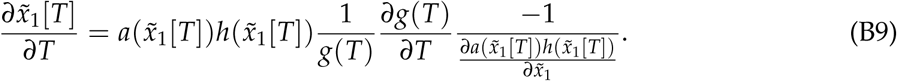

Because 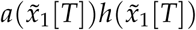 is always positive, then if 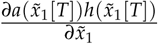 is negative (resp. positive) the sensitivity of 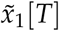 to changes in T is similar (resp. opposite) to that of *g*(*T*) to changes in T (i.e. 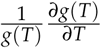).

The sensitivity of 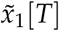 to changes in T depends on the sign of 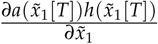.

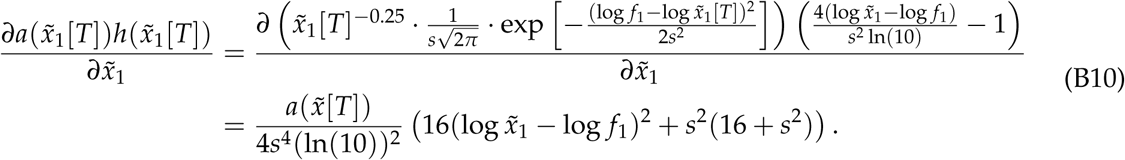

Because 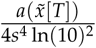 is always positive, we focus on finding the roots of 16(Y — log *f*_1_)^2^ + *s*^2^ (16 + *s*^2^), with 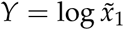. This polynomial of degree 2 is negative (meaning similar impact of temperature on attack rate and consumer body mass) if and only if:

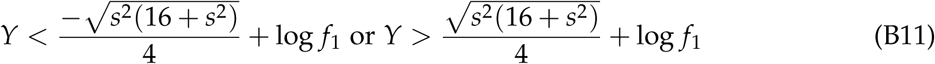

Whether temperature affects consumer body mass similarly to how it affects consumer attack rate thus depends on the relative values of consumer body mass at the singular strategy, degree of generalism s and feeding center *f*_1_. For the typical values used in this analysis, the consumer body mass values at the singular strategy are always above the larger root 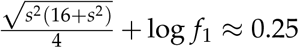. This is coherent with our results on fig. 3A, C and D where the relation between body mass and temperature follows the same direction than the relation between attack rate and temperature. More generally, we expect consumer body masses to be often above the bigger root. Indeed, all consumers in our model have by definition a larger body mass than the external resource, meaning that at evolutionary equilibrium 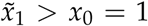, and therefore, *Y* > 0. Values below the larger root would require a high degree of generalism *s* and a consumer with above-optimum feeding center (*f*_1_ > *x*_0_ = 1), such values are thus unlikely.

#### Evolutionary stability analysis - Invasibility

Study of the singular strategy invasibility (i.e. evolutionary stability) requires the second derivative of the fitness with respect to *x_m_* (Marrow et al., 1996). The singular strategy is evolutionarily stable if 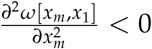. We find that

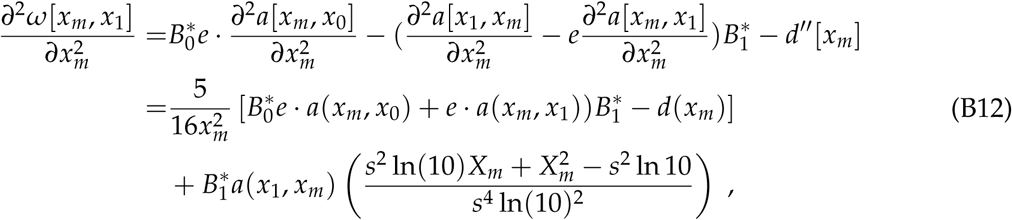

with *X_m_* = log *x_m_* — log *f*_1_, which simplifies into:

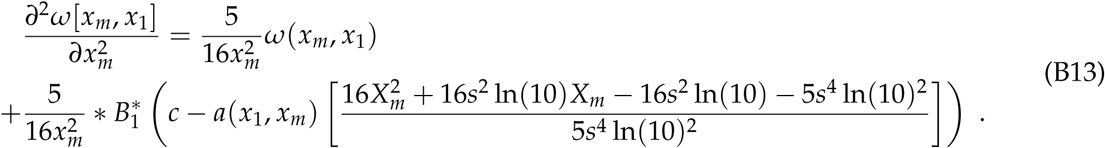

We know that *ω*[*x_m_*,*x*_1_] → 0 in case *x_m_* → *x*_1_. In consequence, we have:

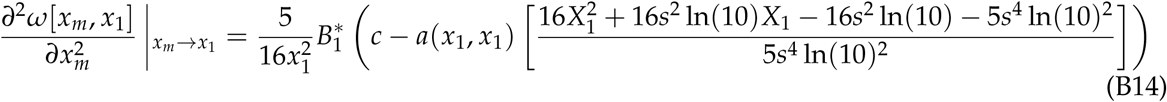

with *X*_1_ = log *x*_1_ — log *f*_1_.

Note that 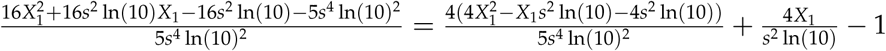.

Therefore, using equation (B5), we can write:

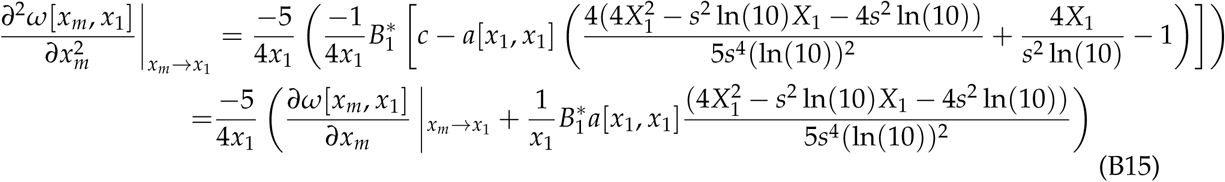

When 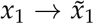 (the predator body mass reaches the singular strategy), then by 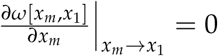 definition. In summary, we find:

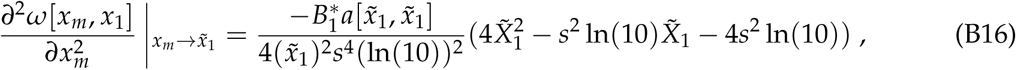

with 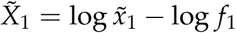.

Since 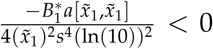, the singular strategy is evolutionarily stable if and only if we have 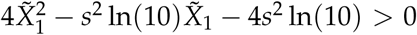. Because the value of 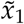 is fixed by equation (15), the sign of equation (B16) will depend on the values of *f*_1_ and *s* that describe the predator feeding niche. Results of such parameter variation are mentioned in the next part on convergence stability. With the typical parameters used in the main article (*f*_1_ = 1 and *s* = 0.25), the bigger singular strategies 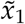 is approximately equal to 6.115 for the standard temperature *T_ref_* = 293K and 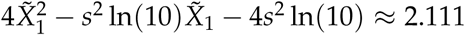. It ensures non-invasibility of this strategy, in consistency with our simulation results.

#### Evolutionary stability analysis - Convergence

Convergence stability conditions are normally computed via either the sum of two second partial derivatives or the derivative of the selection gradient. The singular strategy is convergent stable if:

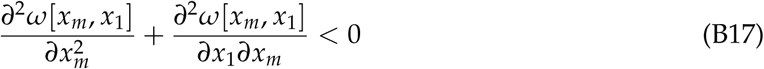

Formula of the first term of this equation is given by equation (B12). The calculation of the cross-derivative is, however, more complex. It corresponds to:

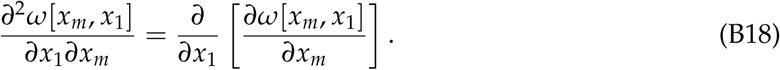

Using the expression of the first derivative 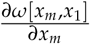 from equation (B4) gives:

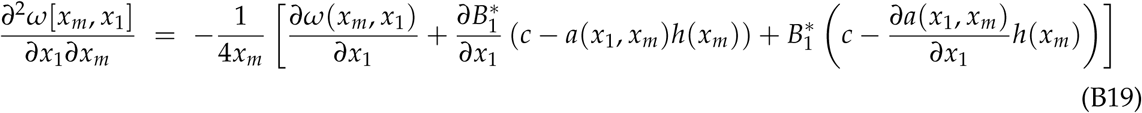

with 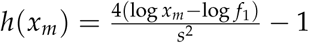 as in equation (B7), and 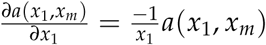. Derivative of the fitness function with respect to *x*_1_ gives:

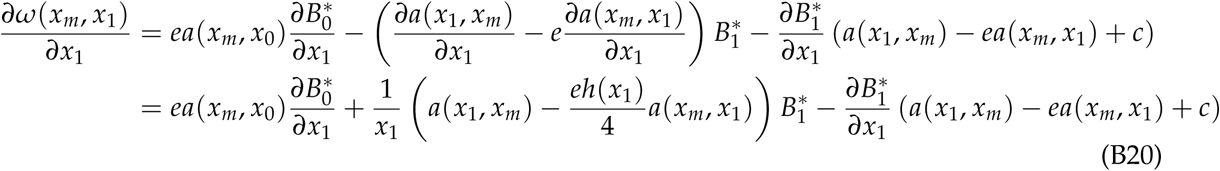

Derivative of 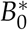 with respect to *x*_1_ gives:

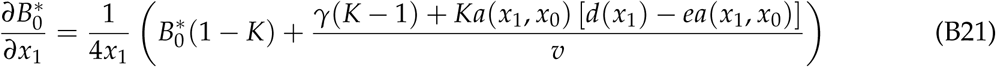

Derivative of 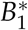 with respect to *x*_1_ gives:

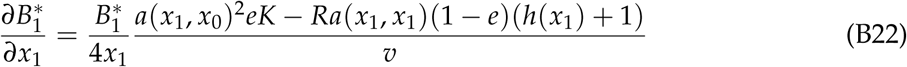

with *γ* = —*R* [*c* — *a*(*x*_1_,*x*_1_)(1 — *e*)*h*(*x*_1_)] and *v* = *a*(*x*_1_,*x*_0_)^2^*eK* + *R* [*a*(*x*_1_,*x*_1_)(1 — *e*) + *c*]. The full cross derivative has then a complex expression. Added to the second derivative, conditions for non-invasibility using equation (B17) are non tractable. Convergence stability of the singular strategy is therefore checked by varying one parameter at a time within the feasibility range. Convergence stability (and evolutionary stability) can then be visually checked by PIPs, and by the calculation of the fitness at 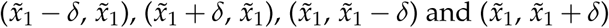 with *δ* = 0.001 kg, around the singular strategy. Convergence stability (resp. evolutionary repelling quality) requires the fitness of 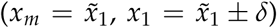 to be positive (resp. negative), because a resident close to the singular strategy can be replaced by a mutant. In this case the mutant reaches the singular strategy, no further mutation is possible. Evolutionary stability requires the fitness of 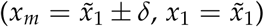 to be negative because a mutant with a slightly bigger or lower body mass than the resident at equilibrium then cannot invade the system.

Parameter variations are given in table B1. For all the parameter values tested in the ranges indicated in table B1, the smallest singular strategy is always a repellor (not convergent stable, not evolutionarily stable), and the biggest always convergent and evolutionarily stable (i.e. a CSS).

**Table B1:** Robustness check for all the model parameters.

| Parameter<sup>1</sup> | Variation range<sup>2</sup> |
| --- | --- |
| $R$ | $[0.5, 10]$ with a 0.5 step |
| $K$ | $[5, 1000]$ with a 5 step |
| $x_0$ | $[0.5, 2.5]^3$ with a 0.05 step |
| $f_1$ | $[0.5, 2.5]^3$ with a 0.05 step |
| $s$ | $[0.05, 1]$ with a 0.05 step |
| $d$ | $[0.05, 1]$ with a 0.05 step |
| $e$ | $[0.3, 1]^3$ with a 0.05 step |
| $c$ | $[0.05, 1]$ with a 0.05 step |

<sup>1</sup> When varying one parameter, all the other are fixed at the typical values specified in table 1.
<sup>2</sup>For all parameter values, we tested all $T$ values in the range $[273\text{K}, \min(T_{lim}, 313\text{K})]$ , with $T_{lim}$ the limit temperature for coexistence, calculated via the formula specified in table A1.
<sup>3</sup> The upper (lower) range value is taken just below (above) the limit value of the parameter for coexistence, with $T = T_{ref}$ .

### C: More information on the multi-trophic model

In this appendix, we present three simulations of the multi-trophic community evolution model in detail, in order to clarify how we collected the data shown in fig. 4 of our main article. The time series of these three simulation runs are shown in fig. C1. They differ only in the temperature (280,290 or 300 K) and in the set of the random numbers. Temperature dependence is included following scenario (a).

**Figure C1:**
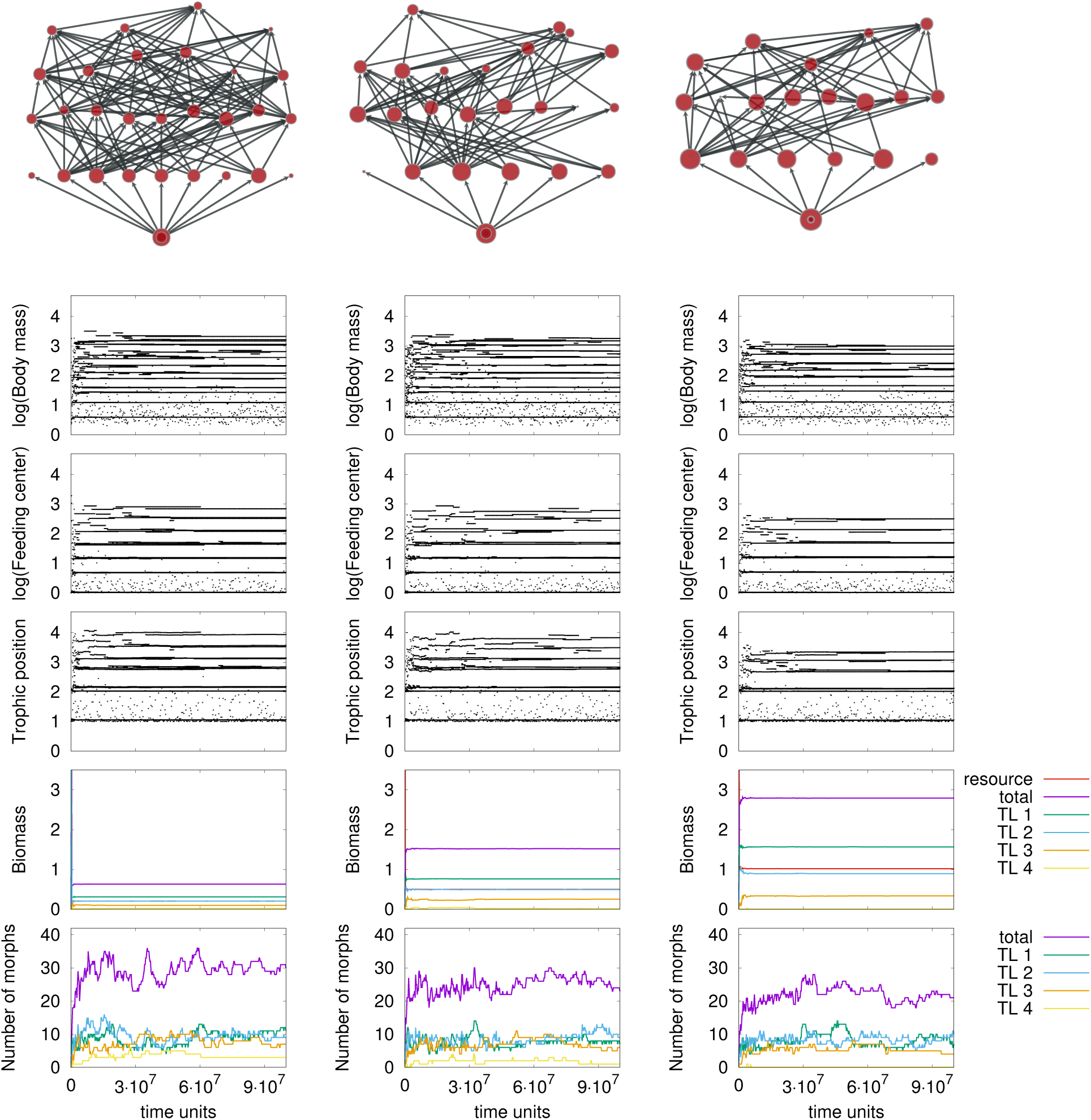
Three exemplary simulation runs of the multi-trophic community evolution model with low (left, *T* = 280*K*), intermediate (middle, *T* = 290*K*) and high (right, *T* = 300*K*) temperature. Temperature dependence is included following scenario (a). The network visualizations at the top of the figure represent the food webs after *t* = 10^8^ time units.

The topmost two panels show the evolution of body masses and feeding centers over time. Each line represents the life span of a morph. Please note that lines might overlap, indicating that several morphs share practically the same trait. Dots represents morphs that emerged, but were not able to establish themselves in the current network. Our mutation rule favors populations with big individual densities, which explains why we generally observe more mutations of morphs with smaller body masses, which typically have the biggest populations.

In the middle panel, we use exactly the same presentation to show the trophic positions of all morphs that are present at a given time. The trophic position of a consumer is calculated as the average, flow-based trophic position of its prey plus one. The trophic position of the external resource is considered to be zero. We round these trophic positions to the next integer value in order to assert all morphs into distinct trophic levels. At a given time, we can now determine how many morphs exist within a given trophic level, as shown in the forth panel, and how much biomass these morphs accumulate, as shown in the fifth panel.

The food webs emerge in a self-organized manner starting with only one single ancestor consumer, which feeds on the external resource. The beginning of the simulations is typically characterized by a period of strong diversification, where higher trophic levels emerge one after the other and where the network structure gets more and more complex. After this initial buildup, we observe that the network size and structure stays approximately the same and only fluctuates around a temperature-dependent average. We are particularly interested in this longterm behavior and therefore deliberately exclude the data from the first 5 · 10^7^ time units from our analysis. Instead, we take only data between *t* = 5 · 10^7^ and the end of the simulation into account. Note that for better clarity fig. C1 shows only the first 10^8^ time steps, although the simulations analyzed for fig. 4 in the main article actually had a much longer runtime of 5 · 10^8^ time units.

## References

Allhoff, K., and B. Drossel. 2016. Biodiversity and ecosystem functioning in evolving food webs. Phil. Trans. R. Soc. B 371:20150281.

Allhoff, K. T., and B. Drossel. 2013. When do evolutionary food web models generate complex structures? J. Theor. Biol. 334:122 - 129.

Allhoff, K. T., D. Ritterskamp, B. C. Rall, B. Drossel, and C. Guill. 2015. Evolutionary food web model based on body masses gives realistic networks with permanent species turnover. Nature Scientific Reports.

Arendt, J. 2007. Ecological correlates of body size in relation to cell size and cell number: patterns in flies, fish, fruits and foliage. Biological Reviews 82:241-256.

Arim, M., F. Bozinovic, and P. A. Marquet. 2007. On the relationship between trophic position, body mass and temperature: reformulating the energy limitation hypothesis. Oikos 116:1524-1530.

Atkinson, D., and R. M. Sibly. 1997. Why are organisms usually bigger in colder environments? making sense of a life history puzzle. Trends in Ecology & Evolution 12:235-239.

Beisner, B. E., E. McCauley, and F. J. Wrona. 1997. The influence of temperature and food chain length on plankton predator prey dynamics. Canadian Journal of Fisheries and Aquatic Sciences 54:586-595.

Bell, G., and A. Gonzalez. 2009. Evolutionary rescue can prevent extinction following environmental change. Ecology letters 12:942-948.

Bergmann, C. 1848. Über die Verhältnisse der Warmeokonomie der Thiere zu ihrer Größe.

Binzer, A., C. Guill, U. Brose, and B. C. Rall. 2012. The dynamics of food chains under climate change and nutrient enrichment. Philosophical Transactions of the Royal Society B: Biological Sciences 367:2935-2944.

Binzer, A., C. Guill, B. C. Rall, and U. Brose. 2016. Interactive effects of warming, eutrophication and size structure: impacts on biodiversity and food-web structure. Global Change Biology 22:220-227.

Bolchoun, L., B. Drossel, and K. T. Allhoff. 2017. Spatial topologies affect local food web structure and diversity in evolutionary metacommunities. Scientific Reports 7.

Brännström, Å., J. Johansson, N. Loeuille, N. Kristensen, T. A. Troost, R. H. R. Lambers, and U. Dieckmann. 2012. Modelling the ecology and evolution of communities: a review of past achievements, current efforts, and future promises. Evolutionary Ecology Research 14:601-625.

Brose, U., J. A. Dunne, J. M. Montoya, O. L. Petchey, F. D. Schneider, and U. Jacob. 2012. Climate change in size-structured ecosystems. Phil. Trans. R. Soc. B 367:2903-2912.

Brose, U., T. Jonsson, E. L. Berlow, P. Warren, C. Banasek-Richter, L.-F. Bersier, J. L. Blanchard, T. Brey, S. R. Carpenter, M.-F. C. Blandenier, et al. 2006a. Consumer-resource body-size relationships in natural food webs. Ecology 87:2411-2417.

Brose, U., R. J. Williams, and N. D. Martinez. 2006b. Allometric scaling enhances stability in complex food webs. Ecol. Lett. 9:1228-1236.

Brown, J. H., J. F. Gillooly, A. P. Allen, V. M. Savage, and G. B. West. 2004. Toward a metabolic theory of ecology. Ecology 85:1771-1789.

Calcagno, V., P. Jarne, M. Loreau, N. Mouquet, and P. David. 2017. Diversity spurs diversification in ecological communities 8:15810.

Calcagno, V., F. Massol, N. Mouquet, P. Jarne, and P. David. 2011. Constraints on food chain length arising from regional metacommunity dynamics. Proceedings of the Royal Society B: Biological Sciences 278:3042-3049.

Cohen, J. E., T. Jonsson, and S. R. Carpenter. 2003. Ecological community description using the food web, species abundance, and body size. Proceedings of the National Academy of Sciences 100:1781-1786.

Daufresne, M., K. Lengfellner, and U. Sommer. 2009. Global warming benefits the small in aquatic ecosystems. Proceedings of the National Academy of Sciences 106:12788-12793.

Dieckmann, U., and R. Law. 1996. The dynamical theory of coevolution: a derivation from stochastic ecological processes. Journal of Mathematical Biology 34:579-612.

Downing, A. L., and M. A. Leibold. 2002. Ecosystem consequences of species richness and composition in pond food webs. Nature 416:837-841.

Dreisig, H. 1981. The rate of predation and its temperature dependence in a tiger beetle, cicindela hybrida. Oikos 36:196-202.

Drossel, B., and A. J. McKane. 2005. Modelling food webs, in Handbook of Graphs and Networks: From the Genome to the Internet, chap. 10. Wiley-VCH Verlag GmbH & Co. KGaA, Weinheim.

Edeline, E., G. Lacroix, C. Delire, N. Poulet, and S. Legendre. 2013. Ecological emergence of thermal clines in body size. Global change biology 19:3062-3068.

Emmrich, M., S. Pédron, S. Brucet, I. J. Winfield, E. Jeppesen, P. Volta, C. Argillier, T. L. Lauridsen, K. Holmgren, T. Hesthagen, et al. 2014. Geographical patterns in the body-size structure of european lake fish assemblages along abiotic and biotic gradients. Journal of Biogeography 41:2221-2233.

Englund, G., G. Öhlund, C. L. Hein, and S. Diehl. 2011. Temperature dependence of the functional response. Ecology letters 14:914-921.

Fedorenko, A. Y. 1975. Feeding characteristics and predation impact of chaoborus (diptera, chao-boridae) larvae in a small lake. Limnology and Oceanography 20:250-258.

Ferriere, R., and S. Legendre. 2013. Eco-evolutionary feedbacks, adaptive dynamics and evolutionary rescue theory. Phil. Trans. R. Soc. B 368:20120081.

Forster, J., A. G. Hirst, and D. Atkinson. 2012. Warming-induced reductions in body size are greater in aquatic than terrestrial species. Proceedings of the National Academy of Sciences 109:19310-19314.

Fussmann, G., M. Loreau, and P. Abrams. 2007. Eco-evolutionary dynamics of communities and ecosystems. Functional Ecology 21:465-477.

Fussmann, K. E., F. Schwarzmüller, U. Brose, A. Jousset, and B. C. Rall. 2014. Ecological stability in response to warming. Nature Climate Change 4:206-210.

Gardner, J. L., A. Peters, M. R. Kearney, L. Joseph, and R. Heinsohn. 2011. Declining body size: a third universal response to warming? Trends in ecology & evolution 26:285-291.

Geritz, S. a. H., E. Kisdi, G. Meszena, and J. a. J. Metz. 1998. Evolutionarily singular strategies and the adaptive growth and branching of the evolutionary tree. Evolutionary Ecology 12:35-57.

Gibert, J. P., and J. P. DeLong. 2014. Temperature alters food web body-size structure. Biology Letters 10.

Gilbert, B., T. D. Tunney, K. S. McCann, J. P. DeLong, D. A. Vasseur, V. Savage, J. B. Shurin, A. I. Dell, B. T. Barton, C. D. Harley, et al. 2014. A bioenergetic framework for the temperature dependence of trophic interactions. Ecology letters 17:902-914.

Gillooly, J. F., J. H. Brown, G. B. West, V. M. Savage, and E. L. Charnov. 2001. Effects of size and temperature on metabolic rate. science 293:2248-2251.

Gonzalez, A., O. Ronce, R. Ferriere, and M. E. Hochberg. 2013. Evolutionary rescue: an emerging focus at the intersection between ecology and evolution.

Gresens, S. E., M. L. Cothran, and J. H. Thorp. 1982. The influence of temperature on the functional response of the dragonfly celithemis fasciata (odonata: Libellulidae). Oecologia 53:281-284.

Hessen, D. O., M. Daufresne, and H. P. Leinaas. 2013. Temperature-size relations from the cellular-genomic perspective. Biological Reviews 88:476-489.

Hulot, F. D., G. Lacroix, F. Lescher-Moutoue, and M. Loreau. 2000. Functional diversity governs ecosystem response to nutrient enrichment. Nature 405:340.

Kirkpatrick, M., and N. H. Barton. 1997. Evolution of a species’ range. The American Naturalist 150:1-23.

Kokko, H., A. Chaturvedi, D. Croll, M. C. Fischer, F. Guillaume, S. Karrenberg, B. Kerr, G. Rolshausen, and J. Stapley. 2017. Can evolution supply what ecology demands? Trends in Ecology & Evolution.

Kozłowski, J., M. Czarnołeski, and M. Dańko. 2004. Can optimal resource allocation models explain why ectotherms grow larger in cold? Integrative and Comparative Biology 44:480-493.

Lavergne, S., N. Mouquet, W. Thuiller, and O. Ronce. 2010. Biodiversity and climate change: integrating evolutionary and ecological responses of species and communities. Annual Review of Ecology, Evolution, and Systematics 41:321-350.

Leibold, M. A., M. Holyoak, N. Mouquet, P. Amarasekare, J. Chase, M. Hoopes, R. Holt, J. Shurin, R. Law, D. Tilman, et al. 2004. The metacommunity concept: a framework for multi-scale community ecology. Ecology letters 7:601-613.

Lindstedt, S. L., B. J. Miller, and S. W. Buskirk. 1986. Home range, time, and body size in mammals. Ecology 67:413-418.

Loeuille, N., and M. Loreau. 2005. Evolutionary emergence of size-structured food webs. PNAS 102:5761-5766.

Loeuille, N., and M. Loreau. 2006. Evolution of body size in food webs: does the energetic equivalence rule hold? Ecology Letters 9:171-178.

Macarthur, R., and R. Levins. 1967. The limiting similarity, convergence, and divergence of coexisting species. The American Naturalist 101:377-385.

Marrow, P., U. Dieckmann, and R. Law. 1996. Evolutionary dynamics of predator-prey systems: an ecological perspective. Journal of Mathematical Biology 34:556-578.

Meehan, T. D. 2006. Energy use and animal abundance in litter and soil communities. Ecology 87:1650-1658.

Metz, J. A. J., R. M. Nisbet, and S. A. H. Geritz. 1992. How should we define ‘fitness’ for general ecological scenarios? Trends in Ecology & Evolution 7:198-202.

Mohaghegh, J., P. De Clercq, and L. Tirry. 2001. Functional response of the predators podisus maculiventris (say) and podisus nigrispinus (dallas) (het., pentatomidae) to the beet army-worm, spodoptera exigua (hubner) (lep., noctuidae): effect of temperature. Journal of Applied Entomology 125:131-134.

Norberg, J., M. C. Urban, M. Vellend, C. A. Klausmeier, and N. Loeuille. 2012. Eco-evolutionary responses of biodiversity to climate change. Nature Climate Change 2:747-751.

O’Gorman, E. J., L. Zhao, D. E. Pichler, G. Adams, N. Friberg, B. C. Rall, S. Alex, H. Zhang, D. C. Reuman, and G. Woodward. 2017. Unexpected changes in community size structure in a natural warming experiment. Nature Climate Change 7:659-663.

Ohlberger, J. 2013. Climate warming and ectotherm body size-from individual physiology to community ecology. Functional Ecology 27:991-1001.

Oksanen, L., S. D. Fretwell, J. Arruda, and P. Niemela. 1981. Exploitation ecosystems in gradients of primary productivity. The American Naturalist 118:pp. 240-261.

Osmond, M. M., M. A. Barbour, J. R. Bernhardt, M. W. Pennell, J. M. Sunday, and M. I. O’Connor. 2017. Warming-induced changes to body size stabilize consumer-resource dynamics. The American Naturalist 189:000-000.

Peck, Y. S. 2016. A Cold Limit to Adaptation in the Sea. Trends in Ecology & Evolution 31:13-26.

Perrin, N. 1995. About berrigan and charnov’s life-history puzzle. Oikos 73:137-139.

Petchey, O. L., A. P. Beckerman, J. O. Riede, and P. H. Warren. 2008. Size, foraging, and food web structure. Proceedings of the National Academy of Sciences 105:4191-4196.

Petchey, O. L., P. T. McPhearson, T. M. Casey, and P. J. Morin. 1999. Environmental warming alters food-web structure and ecosystem function. Nature 402:69.

Peters, R. H. 1986. The ecological implications of body size, vol. 2. Cambridge University Press.

Pillai, P., A. Gonzalez, and M. Loreau. 2011. Metacommunity theory explains the emergence of food web complexity. Proceedings of the National Academy of Sciences 108:19293-19298.

Pimm, S. L. 1982. Food webs. Springer Netherlands.

Pörtner, H. O., and R. Knust. 2007. Climate change affects marine fishes through the oxygen limitation of thermal tolerance. science 315:95-97.

Post, D. M. 2002. The long and short of food-chain length. Trends in Ecology & Evolution 17:269-277.

Rall, B. C., U. Brose, M. Hartvig, G. Kalinkat, F. Schwarzmüller, O. Vucic-Pestic, and O. L. Petchey. 2012. Universal temperature and body-mass scaling of feeding rates. Phil. Trans. R. Soc. B 367:2923-2934.

Rall, B. C., O. Vuvic-Pestic, R. B. Ehnes, M. Emmerson, and U. Brose. 2010. Temperature, predator-prey interaction strength and population stability. Global Change Biology 16:2145-2157.

Reading, C. J. 2007. Linking global warming to amphibian declines through its effects on female body condition and survivorship. Oecologia 151:125-131.

Riede, J. O., B. C. Rall, C. Banasek-Richter, S. A. Navarrete, E. A. Wieters, M. C. Emmerson, U. Jacob, and U. Brose. 2010. Scaling of food-web properties with diversity and complexity across ecosystems. Advances In Ecological Research 42:139-170.

Sentis, A., A. Binzer, and D. S. Boukal. 2017. Temperature-size responses alter food chain persistence across environmental gradients. Ecology Letters.

Shelomi, M. 2012. Where are we now? bergmann’s rule sensu lato in insects. The American Naturalist 180:511-519.

Sheridan, J. A., and D. Bickford. 2011. Shrinking body size as an ecological response to climate change. Nature climate change 1:401.

Sinervo, B., F. Méndez-de-la Cruz, D. B. Miles, B. Heulin, E. Bastiaans, M. V.-S. Cruz, R. Lara-Resendiz, N. Martińez-Méndez, M. L. Calderón-Espinosa, R. N. Meza-Lázaro, H. Gadsden, L. J. Avila, M. Morando, I. J. D. l. Riva, P. V. Sepulveda, C. F. D. Rocha, N. Ibargüengoytía, C. A. Puntriano, M. Massot, V. Lepetz, T. A. Oksanen, D. G. Chapple, A. M. Bauer, W. R. Branch, J. Clobert, and J. W. Sites. 2010. Erosion of Lizard Diversity by Climate Change and Altered Thermal Niches. Science 328:894-899.

Stegen, J. C., R. Ferriere, and B. J. Enquist. 2011. Evolving ecological networks and the emergence of biodiversity patterns across temperature gradients. Proceedings of the Royal Society B: Biological Sciences page rspb20111733.

Tansey, M. R., and T. D. Brock. 1972. The Upper Temperature Limit for Eukaryotic Organisms. Proceedings of the National Academy of Sciences of the United States of America 69:2426-2428.

Thomas, C. D., A. Cameron, R. E. Green, M. Bakkenes, L. J. Beaumont, Y. C. Collingham, B. F. Erasmus, M. F. De Siqueira, A. Grainger, L. Hannah, et al. 2004. Extinction risk from climate change. Nature 427:145-148.

Thompson, D. J. 1978. Towards a realistic predator-prey model: The effect of temperature on the functional response and life history of larvae of the damselfly, ischnura elegans. Journal of Animal Ecology 47:757-767.

Tylianakis, J. M., R. K. Didham, J. Bascompte, and D. A. Wardle. 2008. Global change and species interactions in terrestrial ecosystems. Ecology letters 11:1351-1363.

Urban, M. C., L. De Meester, M. Vellend, R. Stoks, and J. Vanoverbeke. 2012. A crucial step toward realism: responses to climate change from an evolving metacommunity perspective. Evolutionary Applications 5:154-167.

Uszko, W., S. Diehl, G. Englund, and P. Amarasekare. 2017. Effects of warming on predator-prey interactions-a resource-based approach and a theoretical synthesis. Ecology Letters.

van der Have, T. M., and G. de Jong. 1996. Adult size in ectotherms: Temperature effects on growth and differentiation. Journal of Theoretical Biology 183:329-340.

Vasseur, D. A., and K. S. McCann. 2005. A mechanistic approach for modeling temperature-dependent consumer-resource dynamics. The American Naturalist 166:184-198.

Voigt, W., J. Perner, A. J. Davis, T. Eggers, J. Schumacher, R. Bährmann, B. Fabian, W. Heinrich, G. Köhler, D. Lichter, et al. 2003. Trophic levels are differentially sensitive to climate. Ecology 84:2444-2453.

Walther, G.-R. 2010. Community and ecosystem responses to recent climate change. Philosophical Transactions of the Royal Society B: Biological Sciences 365:2019-2024.

West, D. C., and D. M. Post. 2016. Impacts of warming revealed by linking resource growth rates with consumer functional responses. Journal of Animal Ecology.

Willis, K., and G. MacDonald. 2011. Long-term ecological records and their relevance to climate change predictions for a warmer world. Annual Review of Ecology, Evolution, and Systematics 42:267-287.

Woodward, G., B. Ebenman, M. Emmerson, J. M. Montoya, J. M. Olesen, A. Valido, and P. H. Warren. 2005. Body size in ecological networks. TRENDS in Ecology and Evolution 20:403.

Yodzis, P., and S. Innes. 1992. Body size and consumer-resource dynamics. The American Naturalist pages 1151-1175.

Yom-Tov, Y., N. Leader, S. Yom-Tov, and H. J. Baagøe. 2010. Temperature trends and recent decline in body size of the stone marten Martes foina in Denmark. Mammalian Biology - Zeitschrift für Saugetierkunde 75:146-150.

Yom-Tov, Y., and J. Yom-Tov. 2005. Global warming, Bergmann’s rule and body size in the masked shrew Sorex cinereus Kerr in Alaska. Journal of Animal Ecology 74:803-808.

Yom-Tov, Y., S. Yom-Tov, J. Wright, C. J. R. Thorne, and R. Du Feu. 2006. Recent changes in body weight and wing length among some British passerine birds. Oikos 112:91-101.

Yvon-Durocher, G., J. M. Montoya, M. Trimmer, and G. Woodward. 2011. Warming alters the size spectrum and shifts the distribution of biomass in freshwater ecosystems. Global Change Biology 17:1681-1694.

